# Neural Sensitivity to Natural Image Statistics Changes during Middle Childhood

**DOI:** 10.1101/2020.03.23.003905

**Authors:** Benjamin Balas, Alyson Saville

## Abstract

Natural images have lawful statistical properties that the adult visual system is sensitive to, both in terms of behavior and neural responses to natural images. The developmental trajectory of sensitivity to natural image statistics remains unclear, however. In behavioral tasks, children appear to slowly acquire adult-like sensitivity to natural image statistics during middle childhood (Ellemberg et al., 2012), but in other tasks, infants exhibit some sensitivity to deviations of natural image structure (Balas & Woods, 2014). Here, we used event-related potentials (ERPs) to examine how sensitivity to natural image statistics changes during childhood at distinct stages of visual processing (the P1 and N1 components). We asked children (5-10 years old) and adults to view natural texture images with either positive/negative contrast, and natural/synthetic texture appearance (Portilla & Simoncelli, 2000) to compare electrophysiological responses to images that did or did not violate natural statistics. We hypothesized that children may only acquire sensitivity to these deviations from natural texture appearance late in middle childhood. Counter to this hypothesis, we observed significant responses to unnatural contrast and texture statistics at the N1 in all age groups. At the P1, however, only young children exhibited sensitivity to contrast polarity. The latter effect suggests greater sensitivity earlier in development to some violations of natural image statistics. We discuss these results in terms of changing patterns of invariant texture processing during middle childhood and ongoing refinement of the representations supporting natural image perception.

## Introduction

Natural images have lawful properties that the adult visual system is tuned to. For example, the amplitude spectrum of natural scenes in the Fourier domain has been demonstrated to be of the form 1/f^n^, where ‘n’ is approximately 1 (Ruderman & Bialek, 1994; van der Schaaf & van Hateren, 1996), reflecting a systematic fall-off of spectral power from low frequencies to high. The adult visual system exhibits an analogous fall-off in contrast sensitivity from low frequencies to high (Field, 1987), which has been taken as evidence that visual processing is optimized for the spatial content of the natural visual world. Direct tests of this hypothesis have been carried in a variety of ways, most of which have focused on characterizing observers’ sensitivity to patterns that vary according to the exponent that determines the fall-off of spectral power in an image. The basic conjecture is that if adult visual processing is indeed optimally tuned to process images of natural scenes, than sensitivity should be highest for patterns that adhere to the spectral fall-off of natural scenes and lower for patterns that differ from this. This pattern of results was observed using fractal noise images (Knill, Field & Kersten, 1990) and natural scenes (Tadmor & Tolhurst, 1994; Hansen & Hess, 2006), providing support for the tuning of adult vision to natural scene statistics.

Besides examining sensitivity to the slope of natural scene amplitude spectra, tuning to the properties of natural images has also been investigated by using alternative means of disrupting the underlying statistical properties of natural scenes. For example, the application of texture synthesis models to natural images yields synthetic stimuli that are guaranteed to be matched to their parent images within local neighborhoods with regard to a range of features (e.g. linear filter outputs (Heeger & Bergen, 1993) or joint wavelet statistics (Portilla & Simoncell, 2000), but which deviate from those images with regard to higher-order statistics. Those higher-order statistical deviations cannot always be clearly defined in terms of specific computations, but examining the differences between original images and synthetic images generated with or without specific features included in the model (Balas, 2006) suggests that extended contours, global image layout, and coarse-scale feature co-occurrence are generally not captured by existing parametric texture synthesis models. Critically, though there are more recent models that make it possible to match synthetic textures to their parent images more closely with regard to these higher-order structural features, the absence of this matching using simpler models makes it possible to examine how adult visual processing is affected by the absence of these relationships from natural images. In general, removing these statistical regularities from natural images incurs a measurable cost in a range of tasks including invariant texture matching (Balas & Conlin, 2015a) and material categorization (Balas & Schmidt, 2017). Moreover, observers’ ability to distinguish between putative “scene metamers” that are defined by matching parametric texture features like those described above (Freeman & Simoncelli, 2011; Wallis, Bethge & Wichmann, 2016) suggests that the adult visual system is sensitive to the absence of higher-order statistical regularities that are absent from synthetic images rendered via these models. The tuning of human vision to natural scene statistics is thus evident in multiple ways: Observers are sensitive both to relatively simple Fourier-domain properties of natural scenes, and also to higher-order statistical regularities that reflect more complex structures that are present in natural images.

How does sensitivity to natural image statistics develop? To date, there has been relatively little research focused on this question, though there are several reasons to think that visual sensitivity to the regularities described above may emerge slowly. In particular, the developing visual system does not reach adult-like levels of sensitivity until well into middle childhood, even when we consider low-level visual processing. For example, Ellemberg et al. (1999) demonstrated that children’s contrast sensitivity is not fully mature until approximately 7 years of age. More recent results have confirmed that contrast sensitivity continues to develop into the 7-9 year age range (Leat et al., 2009) further supporting the slow development of spatial vision during childhood. Spatial frequency discrimination takes even longer to reach adult-like levels, only reaching maturity at approximately 10 years of age (Ellemberg, Lepore, & Turgeon, 2010). Besides these low-level processes, children’s ability to integrate visual information over large spatial scales also appears to develop slowly, taking well into middle childhood and adolescence to reach adult-like levels of performance. Kovacs (2000) demonstrated, for example, that children’s perceptual organization abilities continue to mature during middle childhood. Contour integration specifically appears to develop slowly (Hipp et al., 2014; Hadad, Maurer & Lewis, 2010), which may also point to a more general failure of spatial integration affecting performance in some tasks positively (children are less sensitive than adults to the Ebbinghaus illusion, (Doherty et al., 2010), while proving sub-optimal in other settings (Jeon et al. 2010). Considered together, these results suggest that children may have reduced sensitivity to the differences between stimuli that match the statistical properties of natural scenes and those that don’t. Whether we consider the specific form of the amplitude spectrum in natural images or the typical incidence of extended contours and coarse-scale structures, children may lack the necessary apparatus to successfully distinguish natural scenes from manipulated images that deviate from those properties. To our knowledge, one of the few studies to directly examine this issue in childhood is Ellemberg et al. (2012), in which children were asked to discriminate images with different amplitude fall-off coefficients centered on values that were either typical of natural scenes, or outside the range of typically observed values. Briefly, while adults and 10-year-old children did not differ in their sensitivity across coefficient values (and were most sensitive to values close to natural image properties), younger children (6- and 8-year-olds) had elevated thresholds relative to adults for those same coefficienet values. This important result suggests that children indeed lack adult-like tuning to natural scene properties, specifically with regard to the amplitude spectrum of natural images.

In the current study, we chose to examine children’s sensitivity to natural image statistics by disrupting two distinct aspects of natural appearance: higher-order texture statistics and contrast polarity. To disrupt the former, we applied a parametric texture synthesis model (Portilla & Simoncelli, 2000) to natural images depicting a range of objects, surfaces, and textures. While the model guarantees that a large family of joint wavelet statistics are matched within local neighborhoods, larger-scale structures including extended contours, extended surfaces, and specularities are typically not preserved. The resulting images are thus well-matched locally to nature images, but substantially different globally and at coarser spatial scales. To disrupt contrast polarity, we negated chroma and luminance values in our parent images. This yielded images that have the same global layout of edges as the original stimuli (sharing *isophotes*, or contours of constant intensity with their parent images – Fleming & Bulthoff, 2015), but locally have opposite gradients of color and luminance everywhere. In both cases, these images are readily perceived as unusual by adult observers, but they differ from natural images via distinct disruptions of statistical regularities that are in both cases distinct from the properties of the amplitude fall-off coefficient discussed above. Our question was whether or not children between the ages of 5-10 years of age would exhibit the same sensitivity to these manipulations as adults, or if children would demonstrate slow development (possibly along different trajectories) of sensitivity to these two disruptions of natural image statistics.

To investigate this question, we chose to use event-related potentials (ERPs) to measure the sensitivity of early visual components to these two manipulations in children and adults. There are several advantages to using ERPs to address this question. First, the use of ERPs makes it possible to side-step some of the challenges associated with coming up with a behavioral task that can be meaningfully performed by both children and adults. For example, simply distinguishing manipulated images from natural scenes is more or less trivial given the two manipulations we described above, while finer-grained questions about image structure might be hard for young children to understand. An ERP paradigm allows us to use a simple attention task to ensure that all observers attend to the stimuli and use neural markers of visual processing as our proxies for sensitivity. Second, ERPs also allow us to examine distinct stages of visual processing by comparing profiles of sensitivity to our stimulus manipulations at different ERP components. Here, we opted to examine both the P1 and the N1, which are temporally distinct components of visual ERPs that reflect early visual processing. Finally, using neural markers of sensitivity to natural image statistics provides a more direct link between our study and ongoing research describing the properties of receptive fields in early stages of the developing visual system (Dekker et al., 2019).

We predicted that compared to adults, young children (5-7 years old) would exhibit reduced sensitivity to the difference between natural and manipulated images at both the P1 and the N1. We expected that older children (8-10 years old) would not differ from adult patterns of sensitivity, in keeping with behavioral data suggesting that low-level visual processing and higher-order spatial integration are both reaching maturity during these years. Both predictions reflect our underlying hypothesis that sensitivity to natural image statistics continues to develop during middle childhood, reflecting ongoing tuning of mechanisms for spatial vision to optimally respond to the structures that are present in natural images.

## Methods

### Participants

Our final sample was comprised of 48 participants: 16 children between the ages of 5-7 years (9 female, Average age = 6;4), 16 children between the ages of 8-10 years (10 female, Average age = 9;1), and 16 adults over the age of 18 (10 female, Average age = 19.3 years). We recruited child participants from the Fargo-Moorhead community and recruited adults from the NDSU Undergraduate Psychology Study Pool. All participants self-reported normal or corrected-to-normal vision and were right-handed as assessed by the Edinburgh Handedness Inventory (Oldfield, 1971). We obtained informed consent from all participants (or their guardians) prior to the beginning of the testing session and children older than 7 years of age also provided written assent to participate.

### Stimuli

The stimuli used in this task were taken from the Flickr Materials Database (https://people.csail.mit.edu/lavanya/fmd.html, Sharan, Rosenhotz & Adelson, 2014), which is a large database containing approximately 100 images depicting 10 different material categories. We selected a set of 48 images that were intended to be diverse in terms of color histograms, glossy vs. matte appearance, roughness vs. smoothness, and the spatial scale of objects and object parts included in each photo. We cropped and resized these images to a common size of 512×512 pixels, and then created alternative versions of each image for use in our full experimental design, including: (1) Contrast-negated images – we created these images by inverting color and luminance channels in each image. (2) Synthetic texture images – we created these images using the Portilla-Simoncelli color texture synthesis algorithm (Liang, Simoncelli & Lei, 2000), which is a parametric model of texture appearance that has been widely used to create synthetic images of natural textures that are matched for a range of joint wavelet statistics (Portila & Simoncelli, 2000). (3) Contrast-negated synthetic texture images – By applying both of the transformations above (texture synthesis followed by contrast negation), we created images that had both negated color and luminance channels and synthetic texture appearance.

### Procedure

During our recording session, we asked participants to carry out an oddball detection task while we recorded continuous EEG from scalp electrodes. We measured EEG from all participants using 64-channel Hydrocel Geodesic Sensor Nets by EGI, connected to an EGI 400 NetAmps amplifier. To ensure that sensor nets fit each participant adequately, we measured each participant’s head circumference and marked the target position of the vertex electrode on each participant’s scalp. Next, each sensor net was soaked in a Potassium Chloride solution for approximately 5 minutes, after which we applied the net to the participant’s scalp. Next, we measured electrode impedances and re-seated individual electrodes and/or applied additional KCl solution to individual electrodes to establish stable impedances below 25kW. During EEG recording, continuous EEG was referenced to the vertex electrode with a sampling rate of 250Hz.

We carried out EEG recording sessions in an electrically-shielded and sound-attentuated chamber. Participants were asked to sit in front a 1024×768 LCD monitor approximately 57cm from the display. We presented our stimuli so that they would subtend approximately 5 degrees of visual angle at this viewing distance, but this likely varied across participants due to differences in participant height across age groups. During the task, we presented participants with our full set of stimulus images in a pseudo-randomized order determined for each participant using routines implemented in EPrime v2.0. Each image was presented for 500ms against a white background, after which time the image disappeared. On ∼10% of trials, we presented participants with an oddball image depicting a cartoon mushroom from the Super Mario Brothers franchise, and asked participants to only make a response to this image, withholding responses to all of our stimulus images. After each image disappeared, we imposed a 500ms interval, followed by an additional intertrial interval sampled randomly from a uniform distribution bounded between 500ms-1500ms. Each image was presented one time during the testing session for a total of 192 trials in the entire task (48 parent images presented once in each of 4 stimulus categories). Participants tended to complete the testing session in about 20 minutes. All stimulus presentation and response collection routines were carried out using custom routines written using EPrime v. 2.0. All EEG recording and event marking routines were carried out using NetStation v. 5.0.

## Results

Our goal was to examine the amplitude and latency of early visual event-related potentials (the P1 and N1) in response to natural texture images with distinct disruptions to natural image statistics. To do so, we will first describe how we obtained ERPs from our continuous EEG data, followed by analyses of both components separately. For each target component, we analyzed stimulus and group factors on ERP responses using 2×2×3 mixed-design ANOVAs with stimulus contrast (positive vs. negative) and stimulus texture (natural vs. synthetic) as within-subject factors and age group (5-7 years, 8-10 years, adults) as a between-subjects factor.

### EEG pre-processing

We carried out pre-processing to obtain event-related potentials from continuous EEG data using the NetStation Tools interface in NetStation v5.0. We began by filtering the data obtained from each participant using a bandpass filter with a lower bound of 0.1Hz and an upper bound of 30Hz. Next, we segmented the continuous data using stimulus onset markers that were inserted online during EEG recording via extensions to EPrime v2.0. These segments began 100ms before stimulus onset and terminated at 1000ms after stimulus onset for a total length of 1100ms. We continued by baseline correcting each of these segments by calculating the average value within the 100ms pre-stimulus onset period, and then subtracting this value from the entire segment. We continued by applying artifact detection algorithms to identify and remove trials contaminated by eye movements, eye blinks and saccades, each of which were determined using simple thresholds of either the raw voltage or the voltage range observed at individual sensors. This was followed by the replacement of any bad channels in the sensor array using NetStation’s bad channel interpolation tool, which uses neighboring channels to replace bad channel data with an estimate drawn from the local neighborhood around the target channel. Finally, we calculated an average ERP for each participant by averaging segments together within subject category at each sensor. This subject average was re-referenced using a global average reference.

### ERP component analysis

We identified sensors of interest and time windows of interest for each target component per age group by inspecting the grand average ERP calculated across subjects in each age range. Specifically, we calculated an average ERP collapsed across all conditions for each age group and used this to visually identify sensors where the P1 and N1 were maximal, and to identify time intervals that included each component, accounting for variability across participants. In terms of sensors of interest, this visual inspection led us to select occipital midline sites to analyze both components in all three age groups, specifically sensors 35, 37 and 39 in the Hydrocel sensor array. Our time windows for each component varied somewhat across age groups: P1 component – 5-7 year-olds, 84ms – 168ms; 8-10 year olds, 84ms – 168ms; adults, 76ms – 104ms. N1 component – 5-7 year-olds, 168ms – 240 ms; 8-10 year-olds, 168ms – 220 ms; adults, 104ms-138ms. In Figure 2, we display grand average ERPs across all conditions for each participant group.

**Fig. 1.**
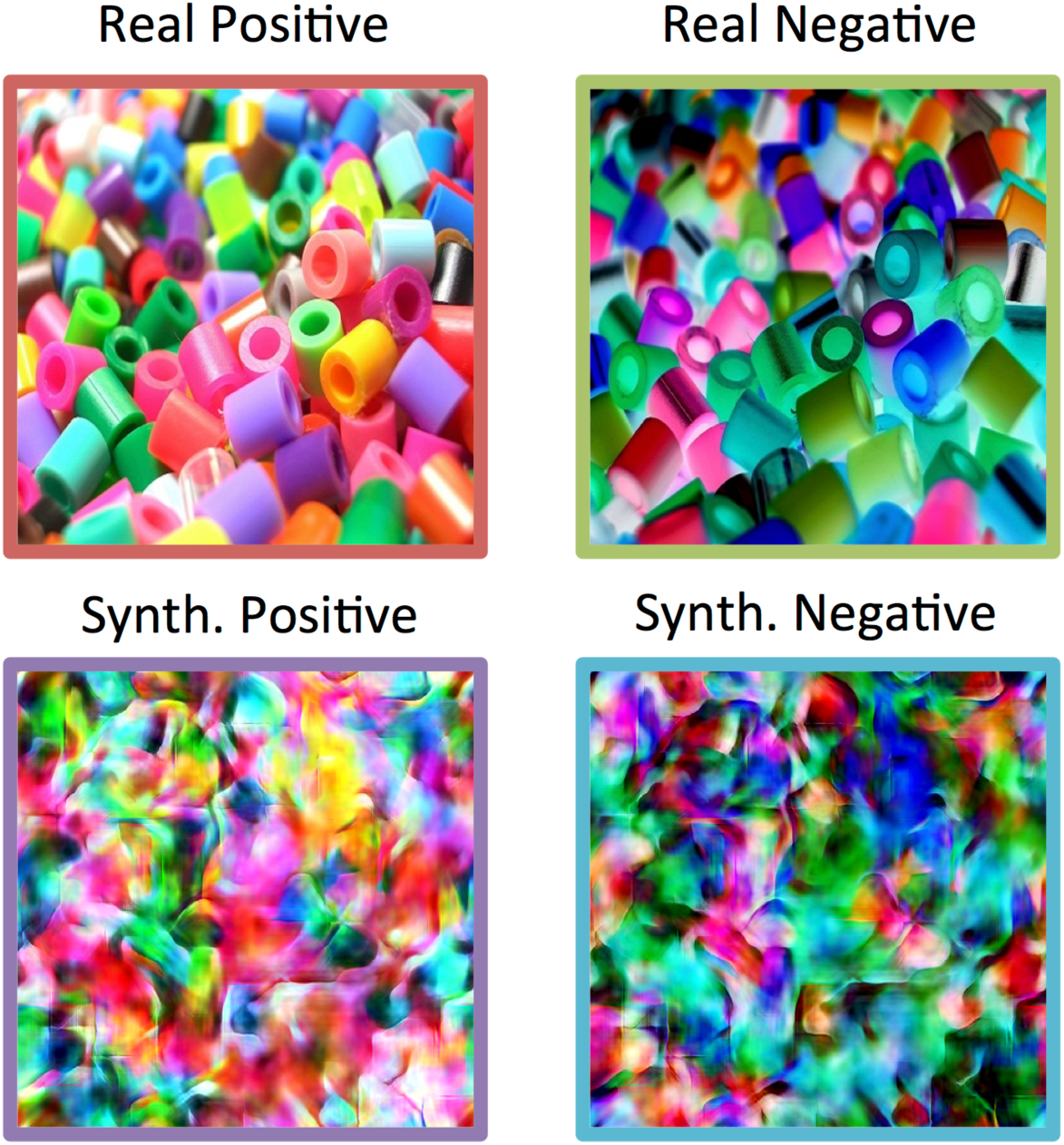
An example image from our task depicting plastic beads in a pile. The original image (Real Positive) was used to create a contrast-negated version (Real Negative), a synthetic texture with the same joint wavelet statistics (Synthetic Positive), and an image to which both manipulations have been applied (Synthetic Negative).

**Fig. 2.**
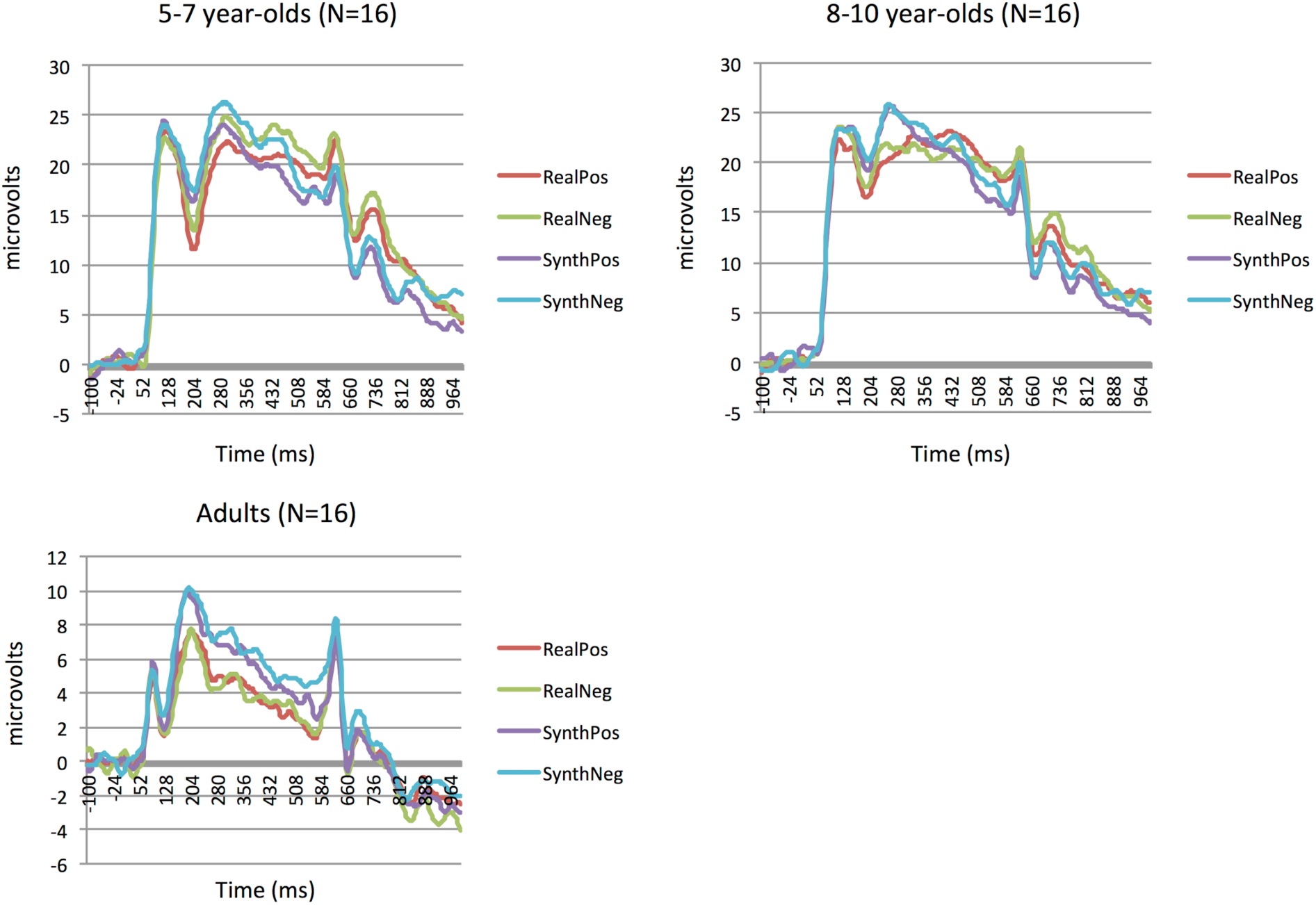
Average ERPs for each stimulus condition for each of our three age groups. These averages were obtained from midline occipital sites in all groups.

Using these sensors and time intervals, we characterized each component using the mean amplitude measured within the time interval for each age group and the peak latency for each component. The extraction of these values across all participants was automated using a Statistic Extraction tool implemented in the NetStation tools interface. We continue by reporting the results of the ANOVAs we carried out using these values for each component. All analyses reported below were carried out using JASP (JASP Team, 2019)

#### P1 mean amplitude

This analysis revealed main effects of stimulus texture (F(1,45)=9.07, p=0.004) and age group (F(2,45)=28.55, p<0.001). The main effect of stimulus contrast did not reach significance (F(1,45)=0.51, p=0.48), nor did we observe any significant interactions between these factors. The main effect of stimulus texture was driven by significantly larger P1 amplitudes in response to synthetic textures compared to real textures (t=-3.01, p_bonf_=0.004). The main effect of age group was the result of significantly larger amplitudes in both child age groups compared to adults (5-7 year-olds vs. adults, t=6,48, p_bonf_<0.001; 8-10 year-olds vs. adults, t=6.6, p_bonf_ <0.001).

#### P1 peak latency

This analysis revealed a main effect of age group (F(2,45)=41.97, p<0.001), but no main effect of either stimulus contrast (F(1,45)=2.78, p=0.10) or stimulus texture (F(1,45)=0.11, p=0.74). The main effect of age group was driven by faster latencies in adults relative to both young children (t=6.6, p_bonf_<0.001) and older children (t=8.8, p_bonf_<0.001) did also observe a significant interaction between stimulus contrast and age group, however (F(2,45)=3.25, p=0.048). No other interactions reached significance. To examine the nature of the interaction between contrast and age, we carried out post-hoc comparisons of the latencies to positive and negative-contrast textures for each age group. This revealed that while young children exhibited significantly longer latencies in response to negative-contrast textures (t=2.16, p_bonf_=0.038), neither older children (t=0.13, p_bonf_=0.90) nor adults (t=0.33, p_bonf_=0.74) exhibited such a difference (Figure 3).

**Figure 3.**
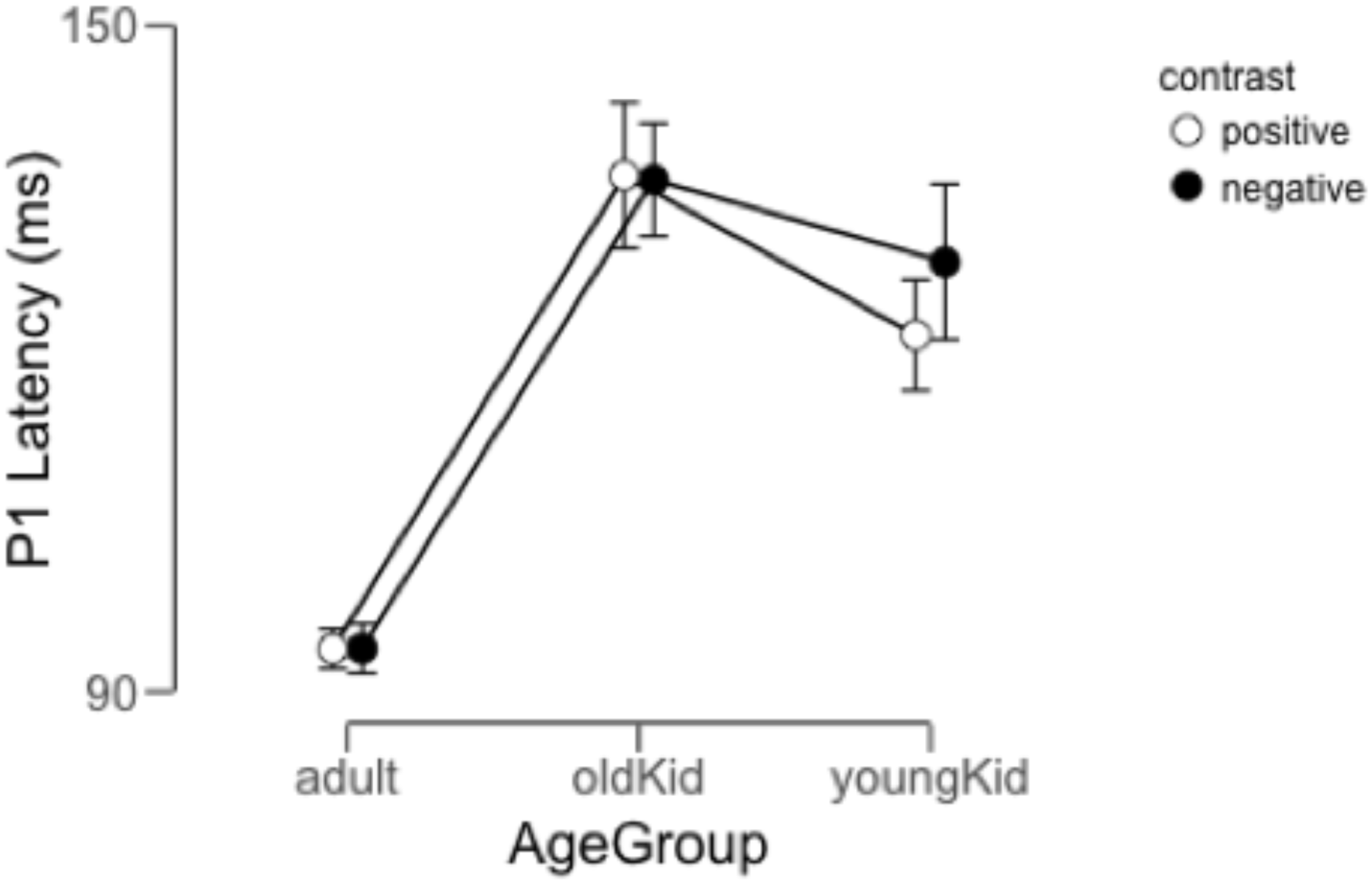
The average P1 latency as a function of age group and contrast. Error bars represent 95% credible intervals calculated via JASP (JASP, 2018).

#### N1 mean amplitude

This analysis revealed main effects of stimulus texture (F(1,45)=35.5, p<0.001), stimulus contrast (F(1,45)=4.72, p=0.035) and age group (F(2,45)=24.5, p<0.001). The main effect of stimulus texture was driven by more negative amplitudes in response to real textures compared to synthetic textures (t=6.2, p_bonf_<0.001), while the main effect of stimulus contrast was driven by more negative amplitudes in response to positive contrast textures compared to negative contrast textures (t=2.37, p_bonf_<0.020). None of the interactions between these factors reached significance. The main effect of age group was the result of significantly more negative amplitudes in both child age groups compared to adults (5-7 year-olds vs. adults, t=5.5, p_bonf_<0.001; 8-10 year-olds vs. adults, t=6.5, p_bonf_ <0.001).

#### N1 peak latency

This analysis revealed only a main effect of participant age (F(2,45)=257.8, p<0.001). Neither the main effect of stimulus contrast (F(1,45)=0.03, p=0.86) nor the main effect of stimulus texture (F(1,45)=0.061, p=0.81) reached significance. The main effect of age group was the result of significantly shorter latencies in both child age groups compared to adults (5-7 year-olds vs. adults, t=20.7, p_bonf_<0.001; 8-10 year-olds vs. adults, t=18.5, p_bonf_ <0.001).We also did not observe any significant interactions between these factors.

## Discussion

Our results demonstrate an intriguing developmental trajectory that suggests differential change during middle childhood with regard to the visual system’s sensitivity to natural image statistics. First, with regard to our adult participants, the data largely replicate previous results we reported using a similar ERP task designed to measure the P1 and N1’s responses to deviations from natural image structure (Balas & Conlin, 2015b). Specifically, our results in both studies indicate that adult participants’ early visual ERP components are sensitive to the losses of extended contours and large-scale image features that follow from applying texture synthesis to a natural image, but are *insensitive* to the difference between positive and negative contrast polarity. Our previous results were based on grayscale stimuli only, and the current study thus extends this result to full-color textures. In some ways, this result is itself somewhat surprising: Why should the profound differences between positive and negative contrast images, which include the reversal of specularities such that they appear as spots, the reversal of luminance edges, and a dramatically remapped color palette, not lead to systematically different responses at early stages of visual processing? We will return to this point later, but for now we simply point out that the adult-like profile of responses favors sensitivity to disruptions of higher-order disruptions of natural image statistics that require extended spatial integration over relatively large image regions to be detected.

Our initial hypothesis was that children’s sensitivity to both manipulations of natural image statistics would be reduced relative to adults, reflecting the ongoing maturation of the underlying mechanisms required to measure those deviations robustly. Instead, our data tell a rather different and perhaps more interesting story. Young children (unlike older children and adults) exhibit sensitivity to contrast polarity as well as sensitivity to the difference between natural and synthesized images. Both of these outcomes violate our expectations in different ways. With regard to children’s sensitivity to natural vs. synthetic image statistics, our data indicate that even though young children do not appear to have mature spatial integration abilities (as evidenced by poorer contour integration, etc. (Jeon et al., 2010; Hadad, Maurer & Lewis, 2010), their visual system is nonetheless sensitive to the absence of higher-order correlations between edge-like features in synthetic texture images. Thus, though the mechanisms we expected might support sensitivity to natural vs. synthetic images are not mature at age 5 (and may not be for years to come (Ellemberg, Hansen & Johnson, 2012)), early stages of the visual system respond differently to our stimuli nonetheless. This may be a function of the magnitude of the difference between our natural and synthetic stimuli with regard to coarse-scale structure, but is still an intriguing result. With regard to contrast polarity, the early sensitivity to positive vs. negative natural images in young children raises an interesting question: Why are young children sensitive to this difference when adults don’t seem to be? There are of course many settings in which negative-contrast images do affect adult performance (see the many studies describing the deleterious effects of contrast negation on face recognition performance, for example (Galper, 1970; Itier & Taylor 2002; Russell et al., 2006)), but in the context of the current study it is clear that early visual ERP components lose sensitivity to contrast polarity in natural scenes during middle childhood. What is the developmental significance of this trajectory?

One potentially interesting way to conceptualize this result is with regard to what has been referred to as “pre-constancy” visual function in young infants (Yang, 2015). Briefly, one aspect of children’s visual development may be acquired invariance (as opposed to sensitivity) to properties of natural images that it is adaptive to produce stable responses to. In the context of object recognition, for example, illumination-invariant recognition requires the observer’s visual system to respond at some level in the same way to images that may differ substantially from one another in terms of where there specific luminance gradients across image regions (Moses, Adini & Ullman, 1997). At early stages of development, before such invariance has been learned, the visual system exhibit a sort of incontinent sensitivity to most image differences, making recognition more challenging due to the ever-changing responses to distinct images of the same objects, faces, surfaces, or textures. With continued development, however, the visual system may learn to produce stable responses to those distinct stimuli and exhibit reduced sensitivity to image differences that previously led to large changes in response. We propose that our results with regard to contrast polarity may reflect a slow version of this developmental trajectory that extends into middle childhood. That is, young children appear to retain sensitivity to local contrast relationships that ultimately they will acquire invariance to, perhaps in service of some high-level visual task that benefits from reduced sensitivity to edge polarity. What tasks might those be? For now, it is difficult for us to say, especially because children’s behavior in tasks involving contrast-negated faces indicates *increasing* sensitivity to contrast polarity with age (Itier & Taylor, 2004). The difference between sensitivity to unnatural appearance in a domain-specific mechanism (e.g., face recognition) and more general visual processing (e.g. natural scene processing), may turn out to be profound.

A further comparison to infant results from a similar paradigm also reveals a more complete developmental trajectory for both types of sensitivity to natural image statistics across the lifespan. Specifically, in previous work with 6-month-old and 9-month-old infants (Balas & Woods, 2014; Balas, Saville & Schmidt, 2018), we found that while infants’ visual ERPs did not exhibit much sensitivity to contrast polarity either early or late in infancy, sensitivity to the difference between natural and synthetic texture appearance was only evident late in infancy. Thus, neural sensitivity to contrast polarity likely emerges after the first year of life, only to disappear during middle childhood. By comparison, sensitivity to synthetic texture appearance emerges sooner (late infancy) and is maintained from then on into adulthood. Again, this trajectory of sensitivity to contrast polarity in the context of natural scene processing differs substantially from what is observed with face-like patterns (Farroni et al., 2005), but still suggests an intriguing pattern of acquired sensitivity and invariance over an extended period of development.

Finally, we close by noting that in all groups, some sensitivity to contrast polarity appeared to be evident at later components of the ERP response, perhaps reflecting the use of contrast polarity information at later stages of visual processing. While these components were not the focus of the current study (and as such, we chose not to offer a post-hoc analysis of their response properties), we speculate that this may be in keeping with our interpretation of reduced sensitivity to contrast polarity as an indicator of an important developmental shift of information towards different stages of processing that support performance in more complex tasks. We suggest that this may be an important area for continued developmental research, as children’s patterns of sensitivity and invariance exhibit intriguing results here with regard to the initial encoding of natural scene structure. Understanding how children’s visual systems are tuned both to the environment in general and also tuned to specific recognition tasks (e.g. face and object recognition) may lead to a deeper understanding how the optimality of the visual system may be defined different criteria at multiple stages of visual processing that nonetheless must work together so that natural stimuli can be efficiently and effectively perceived, distinguished, and recognized.

